# The Effects of Vancomycin on the Viability and Osteogenic Potential of Bone-derived Mesenchymal Stem Cells

**DOI:** 10.1101/316810

**Authors:** Elzaan Booysen, Hanél Sadie-Van Gijsen, Shelly M. Deane, William Ferris, Leon M.T. Dicks

**Affiliations:** Department of Microbiology, Faculty of Natural Sciences, Stellenbosch University, Stellenbosch, South Africa; Division of Endocrinology, Department of Medicine, Faculty of Medicine and Health Sciences, Stellenbosch University Tygerberg Campus, Parow, South Africa

**Keywords:** Vancomycin, mesenchymal stem cells, viability

## Abstract

Periprosthetic joint infections (PJI), caused by methicillin-resistant *Staphylococcus aureus* (MRSA), is the major cause of total hip arthroplasty (THA) failures. Traditionally, MRSA is treated with vancomycin, administrated intravenously or applied directly onto infected tissue. The effect of direct (as opposed to systemic) vancomycin treatment on bone formation and remodelling is largely unknown. The minimal inhibitory concentration (MIC) of vancomycin was determined by adding 200 μl of different concentrations (1 – 20 μg/ml) to actively growing cultures of *S. aureus* Xen 31 (methicillin-resistant) and *S. aureus* Xen 36 (methicillin-sensitive), respectively, and recording changes in optical density over 24 h. Bone marrow-derived and proximal femur-derived mesenchymal stem cells (bmMSCs and pfMSCs) from rat femora were exposed to 1 x MIC (5 μg/ml) and 4 x MIC (20 μg/ml) of vancomycin for 7 days. Cell viability was determined by staining with crystal violet and MTT (3-(4,5- dimethylthiazol-2-yl)-2,5-diphenyltetrazolium bromide), respectively, and osteogenic differentiation by staining with Alizarin Red S. Vancomycin had no effect on the viability of bmMSCs and pfMSCs, even at high levels (20 μg/ml). The osteogenic differentiation of pfMSCs was partially inhibited, while osteogenesis in bmMSCs was not severely affected. The direct application of vancomycin to infected bone tissue, even at excessive levels, may preserve the viability of resident MSC populations. Short-term demineralization may thus be reversed after cessation of vancomycin treatment, improving the outcome of THA surgery.

## INTRODUCTION

The three major load bearing joints in the human body are the ankle, knee and hip (1). The human hip joint has a complex anatomical structure and is responsible for carrying the weight of the upper body. During daily activity, the hip joint may experience loading forces up to 7 times the body weight of the individual, generating immense mechanical pressure which is most likely the main cause for osteoarthritis (OA) in this joint (2). OA effects more than 25% of elderly people and is therefore a major cause of disability in this population group (3, 4).

Currently the only treatment available for severe hip OA is total hip arthroplasty (THA) (5). A metal prosthesis is inserted into the proximal region of the femur, replacing the proximal femur head and neck (6). Although THA is considered the most successful orthopaedic procedure of the past 120 years, and although advances in the field have significantly improved the risk profile associated with this procedure over the past five decades, complications still do occur (7–9). Periprosthetic joint infection (PJI) is the main cause for THA revision surgeries and is defined as bacterial contamination of the joint prosthesis and infection of the adjacent tissue (10). Most, if not all, PJI are caused by bacteria capable of forming biofilms (5). Many strains of these bacteria are resistant to a number of antibiotics (11), and treating PJI is therefore extremely difficult. The most common bacterial infections after THA are Gram-positive cocci. *Staphylococcus aureus* was isolated in 38% of PJI cases. Fifty-three percent of the strains were resistant to methicillin (12).

Failed treatment of PJI with conventional antibiotic regimes places an immense burden on medical professionals. Conventional treatments include intravenous administration of antibiotics for a maximum of 2 weeks, while aggressive treatment includes removal of the infected prosthesis and flushing of the infected joint (one-step revision surgery), followed by another 12 weeks on antibiotics. Once the infection has been cleared, a new prosthesis is inserted (13). In severe cases a surgeon can opt for a two-step revision surgery. This entails initial removal of the infected prosthesis and insertion of an antibiotic-augmented cement spacer, followed by the implantation of a new prosthesis once the bacterial infection has been eradicated (13, 14). Both one-step and two-step revision surgeries can result in prolonged immobility, hospitalization and morbidity. The cost of treatment of a single episode of PJI can escalate to USD 50,000 or more (9). The increase in antibiotic resistance among hospital-acquired infections like PJI has also increased the risk of failing to successfully treat PJI (15, 16). It is therefore clear that PJI presents a major challenge from both a health-care and economic perspective (17).

Methicillin-resistant *Staphylococcus* infections are usually treated with vancomycin (18). Vancomycin is either administered intravenously or directly to the infected site, by means of antibiotic-loaded cement spacers (19–21). However, the effect of direct (as opposed to systemic) vancomycin treatment on bone formation and remodelling is largely unknown. Bone marrow-derived mesenchymal stem cells (bmMSCs) have traditionally been used to study the effects of pharmaceutical treatments on the viability, differentiation and function of osteoblasts (bone-forming cells) *in vitro* (22–24). Recently Jacobs and co-authors (25) reported on a newly characterized MSC, isolated from the proximal end of rat femora. Although these proximal femur MSCs (pfMSCs) are phenotypically similar to bmMSCs, they are functionally distinct, with increased glucocorticoid sensitivity and a reduced osteoblastic differentiation potential in culture. *In vivo*, pfMSCs reside in the exact area of bone tissue that is in contact with the THA prosthesis, and is therefore likely to participate in osseo-integration of the prosthesis. Consequently, these cells can serve as a relevant *in vitro* model to assess the effects of directly applied vancomycin on the MSC populations residing in the bone tissue, which could affect the success of THA revision surgeries. In the present study we evaluate the effects on increasing doses of vancomycin in the viability and osteoblastic differentiation of cultured rat bmMSCs and pfMSCs.

## MATERIALS AND METHODS

### Materials

Bacterial growth medium was from Biolab Diagnostics (Midrand, South Africa), unless stated otherwise. All constituents of osteogenic differentiation media were from Sigma-Aldrich (Schndlldorf, Germany). Dulbecco’s Modified Eagle Medium (DMEM: #BE12-604F; 4.5 g/L glucose), Penicillin/Streptomycin (Pen-Strep: #DE17-602E; 10000 units each in 100 ml), trypsin (#BE17-161E) and Hanks’ Balanced Salt Solution with calcium and magnesium (HBSS: #BE10-508F) were from Lonza (Basel, Switzerland). Fetal bovine serum (FBS) was from Biochrom (Berlin, Germany) and collagenase I (#CLS1) from Worthington Biochemical Corporation (Lakewood, USA). Cell culture dishes were from NEST Biotechnology (New Jersey, USA). Sodium pentobarbitone (Eutha-naze) was from Bayer (Kempton Park, Gauteng, South Africa).

### Determination of MIC (minimum inhibitory concentration)

A micro-broth dilution assay, described by Andrews (26), was used to determine the MIC of vancomycin. *Staphylococcus aureus* Xen 31 (methicillin-resistant *S. aureus*: MRSA) and *S. aureus* Xen 36 (methicillin-sensitive) were used as indicator strains. Both strains were from Bioware™ Microorganisms (Caliper Life Sciences, Hopkinton, USA). Cells from an overnight culture in Brain Heart infusion (BHI) broth, cultured at 37°C, were streaked onto BHI agar and incubated for 24 h at 37°C. A single colony was inoculated into BHI broth and incubated for 18 h at 37°C. Overnight cultures of the two strains were each adjusted to an OD of 0.1 (600 nm). Different concentrations of vancomycin (1 – 20 μg/ml) were added to each liquid culture (final volume of 200 μl) and changes in growth observed by recording absorbance readings at 600 nm, immediately after addition, and 6 h and 24 h later.

### Isolation and maintenance of MSC

Ethical clearance to conduct experiments involving animals was granted by the Research Ethics Committee: Animal Care and Use of Stellenbosch University (reference number SU-ACUD15-00012). Male Wistar rats, 12 weeks old, with an average body mass of 250 g, were housed at the Stellenbosch University Animal Facility and kept according to the guidelines of the South African Medical Research Council. The animals were fed *ad libitum* on standard laboratory feed and sacrificed with an intraperitoneal injection of sodium pentobarbitone (12 mg/kg body mass).

Isolation of bmMSCs and pfMSCs was performed as described by Jacobs and co-workers (25). Femora were surgically removed and cleaned from muscle tissue using sterile gauze. Proximal regions of the femora were removed with sterile surgical side cutters, cut into 1 mm^3^ fragments and digested in Hanks’ Balanced Salt Solution containing 0.075% (w/v) collagenase I and 1.5% (w/v) bovine serum albumin for 1 h at 37°C. Denuded bone fragments were washed five times with DMEM, seeded into a culture dish with isolation media (DMEM containing 1%, v/v, Pen-Strep, and 20%, v/v, FBS) and incubated at 37°C for 24h. Bone fragments were then washed with sterile phosphate-buffered saline (PBS), retained in the dish and submerged in standard growth media (SGM: DMEM containing 1%, v/v, Pen-Strep and 10%, v/v, FBS) for 7 to 10 days to allow migration of pfMSC from the fragments. bmMSCs were flushed from bone marrow cavities with 9 ml (3 ml per flush) cell isolation media and collected in a cell culture dish (100 mm diameter). pfMSCs and bmMSCs were cultured at 37°C for 24 h in 95% humidified air and in the presence of 5% CO_2_. Sterile PBS at 37°C was used to remove non-adherent tissue and the media replaced with SGM.

Both cell types (pfMSCs and bmMSCs) were cultured to 80% confluence and then disaggregated with 1 ml 0.5% (w/v) trypsin and sub-cultured at a ratio of 1:4. All cell cultures were expanded to passage 3 before being used for further experiments. Cell growth media, including media containing supplements, were replaced every 2-3 days. Cells were maintained at 37°C in 95% humidified air containing 5% CO_2_.

### Cytotoxicity of vancomycin

Isolated MSCs at passage 3 were seeded into 12-well plates for crystal violet staining or 96-well plates for the MTT conversion assay and grown until post-confluence in SGM. A combination of cycloheximide (10 μg/ml) and tumour necrosis factor-α (TNF-α; 5 ng/ml) was used as the positive control for cytotoxicity in bmMSCs, while 1 μM dexamethasone (Dex) was used as the positive control for cytotoxicity in pfMSCs, as reported previously (25). Before crystal violet staining, cells were treated with 1 x MIC and 4 x MIC vancomycin, respectively, in SGM for 7 days, fixed with 70% (v/v) ethanol, stained for 5 min with 500 μl 0.01% (w/v) crystal violet solution and rinsed three times with PBS. In these experiments MIC levels were calculated based on results obtained with MRSA strain Xen 31. Bound crystal violet was extracted with 500 μl 75% (v/v) ethanol and absorbance measured at 570 nm. For the MTT conversion assays, cells were treated as described for crystal violet staining, with a final volume of PBS being 100 μl per well. The methodology for the assay was adapted from the protocol for the Sigma-Aldrich *in vitro* Toxicology Assay Kit (MTT-based, #Tox1). After 7 days of treatment with 1 x MIC and 4 x MIC vancomycin, respectively, 10 μl of 5 mg/ml MTT stock solution (Sigma-Aldrich #M2128), dissolved in DMEM, was added to each well. Plates were incubated for 2h at 37°C in the dark and the colour reaction stopped by adding 100 μl of solubilisation solution (10% Triton X-100 diluted with 0.1 N HCL in anhydrous isopropanol) to each well. Solubilisation was aided by incubating the 96-well plate on a plate shaker and repeated mixing with a pipette. Colour development was quantified spectrophotometrically at 690 nm (background absorbance) and 570 nm, respectively. Background absorbance values were subtracted from the A_570_ values. All experiments were performed with quadruplicate biological repeats and triplicate technical repeats.

### MSC differentiation

Isolated MSCs at passage 3 were plated in 12-well culture plates and cultures in SGM until post-confluence as described elsewhere. Osteoblastic differentiation was induced with osteogenic media (OM: SGM supplemented with 50 μM ascorbic acid, 10 mM β-glycerolphosphate and 10 nM dexamethasone), as described by Jacobs *et al*. (25). In parallel, cells were differentiated in the presence of increasing concentrations of vancomycin. Differentiation was evaluated by staining with Alizarin Red S stain (ARS: Amresco, USA) after 7 days (bmMSC) or 21 days (pfMSC). Before staining with 40mM ARS (pH 4.0 – 4.1), cells were washed with PBS, fixed for 5 min at 25°C with 70% (v/v) ethanol and rinsed twice with sterile water. BmMSCs were stained for 2 to 4 hours, after which the excess stain was removed, and cells washed three times with sterile water, once with sterile PBS and an additional three times with sterile water. Bound stain was extracted using 10% (w/v) cetylpyridinium chloride (CPC) dissolved in 10 mM Na_2_HPO_4_ (pH 7.0) and quantified spectrophotometrically at 562 nm. PfMSCs were stained overnight with ARS and washed, as described for bmMSC. Since pfMSCs have reduced osteogenic potential, staining was quantified using image analysis. Images were captured using an Olympus CKX41 microscope (CKX41, CachN 10x0.25 PhP objective), fitted with a Canon EOS 600D camera at 10x magnification. Four random images (one in each quadrant) were taken of each well, resulting in 12 images per treatment since experiments were performed in triplicate. Images were analysed with ImageJ software (version 1.51 J8) and converted into red-green-blue stacks. Analyses were performed in the green channel. A threshold value of T=90 was used to exclude non-specific background staining and remained unchanged throughout. The percentage area stained was recorded and the average for the 12 images per condition was calculated.

### Statistical analysis

GraphPad Prism (version 5.01) was used for all statistical analyses and data were expressed as average ±SD. One-way ANOVA and Dunnett’s *post hoc* test were used to analyse the data. When P < 0.05, the difference was considered to be statistically significant and indicated with an^*^.

## RESULTS AND DISCUSSION

The increase in antibiotic resistance among hospital-acquired infections such as PJI has escalated the risk of PJI treatment failure (15, 16) and may force clinicians to employ ever more aggressive treatment strategies with currently available antibiotics. In the case of PJI, these strategies often involve direct application of antibiotics such as vancomycin to the infected surgical site (13, 14), but the effects of such treatment on the surrounding bone tissue have not been studied at a cellular level. In particular, detrimental effects of antibiotic treatment on the stem cell population residing in bone may result in poor osseointegration of implanted prostheses, leading to structural failure and impaired function of the reconstructed joint even when the PJI has been successfully eradicated.

Based on micro-broth dilution assays, the MIC of vancomycin against *S. aureus* Xen 31 (MRSA) was 5 μg/ml and 1 μg/ml against Xen 36, suggesting that *S. aureus* Xen 31 has reduced susceptibility for vancomycin (27, 28).

Crystal violet stained all cells (dead and alive), while the MTT conversion assay detected only metabolically active cells. High doses of vancomycin, up to 20 μg/ml (4 x MIC for MRSA), had no cytotoxic effects on bmMSCs (Fig. 1A and Fig. 2A) and pfMSC (Fig. 1B and Fig. 2B), as shown with both assays.

**FIG 1.**
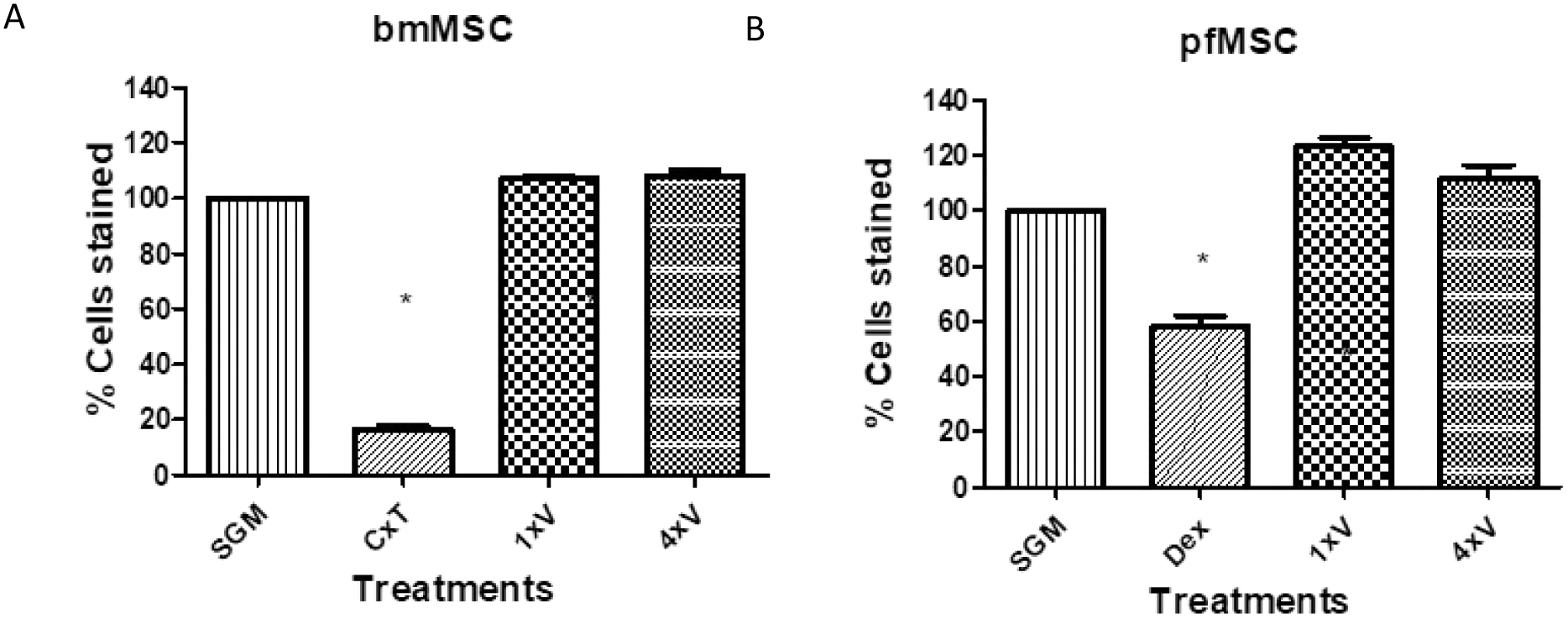
The effect of vancomycin on the cell culture density of bmMSCs (left panel) and pfMSCs (right panel). Cells in SGM were treated for 7 days with 1 x MIC (5 μg/ml) and 4 x MIC (20μg/ml) vancomycin. Positive controls for cytotoxicity were 10 μg/ml cycloheximide together with 5 ng/ml TNFα (CxT) for bmMSCs and 1 μM dexamethasone (Dex) for pfMSCs. The cells were stained with crystal violet. The stain was then extracted and quantified spectrophotometrically at 570 nm. Values for control (SGM-treated) wells were set as 100%. The graph represents data of n = 4. A *post hoc* Dunnet test was done with SGM as control. Statistical significance (p<0.05) is indicated with *.

**FIG 2.**
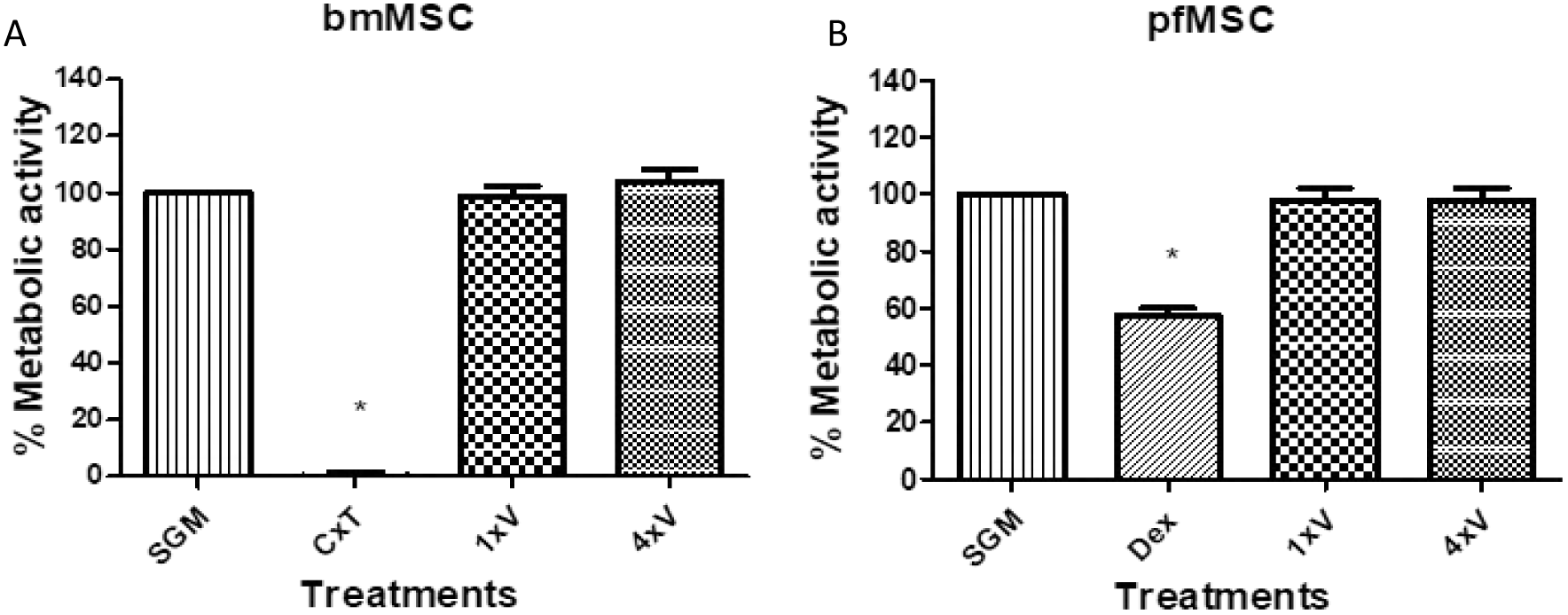
The effect of vancomycin on the metabolic activity of bmMSCs (left panel, A) and pfMSCs (right panel, B). Cells in SGM were treated for 7 days with 1 x MIC (5 μg/ml) and 4 x MIC (20 μg/ml) vancomycin. Positive controls for cytotoxicity were 10 μg/ml cycloheximide and 5 ng/ml TNFα (CxT) for bmMSCs and 1 μM dexamethasone (Dex) for pfMSCs. Cell viability was measured via an MTT conversion assay and intensity of the colour product was measured at 570 nm. Values for control (SGM-treated) cells were set as 100%. The graph represents data of n = 4. A *post hoc* Dunnet test was done with SGM as control. Statistical significance (p < 0.05) is indicated with *.

Based on ARS staining, bmMSCs were fully differentiated after 7 days of treatment with OM, while pfMSCs formed individual mineralized nodules after 21 days of OM treatment. Mineralization in bmMSCs increased slightly, but not significantly, when treated with 4 x MIC vancomycin (Fig. 3A). In contrast, pfMSCs treated with vancomycin showed significantly reduced mineralization, compared to OM-treated cells (Fig. 3B).

**FIG 3.**
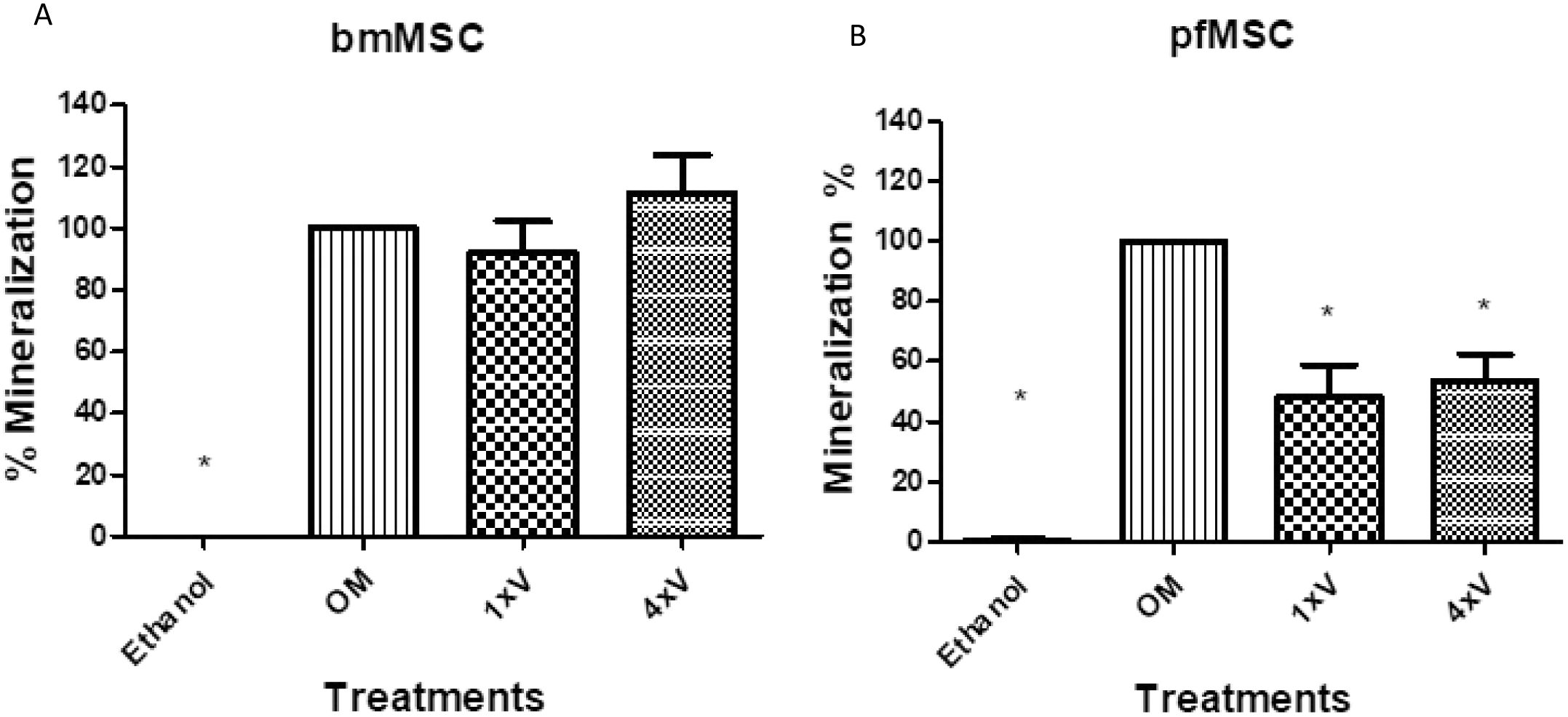
The effect of vancomycin on the osteoblastic differentiation of bmMSC (left panel, A) and pfMSC (right panel, B). Cells were treated with with 1 x MIC (5 μg/ml) and 4 x MIC (20 μg/ml) vancomycin. The formation of mineralized extracellular matrix was evaluated with ARS staining after 7 days (bmMSCs) or 21 days (pfMSCs). Cells treated with control media (SGM plus 0.1%, v/v, ethanol) did not form any mineralized deposits. The graph represents n = 7. All data were compared to OM, which was set as 100%. Statistical significance is indicated with an *.

Vancomycin inhibitions bacterial cell wall synthesis by binding to the terminal D-Ala-D-Ala dipeptide of peptidoglycan units (29). This may explain why vancomycin had no cytotoxic effect on MSCs. However, the mechanisms involved in the reduction of mineralization, specifically in pfMSCs, but not in bmMSCs, remains unclear. The two cell types react differently to external factors, as also shown with exposure to glucocorticoids (25). In the latter study, the authors have shown that pfMSCs are more sensitive to glucocorticoids than bmMSCs.

Our findings are of particular relevance for THA, as pfMSCs reside in the area directly involved in THA (6). Our results suggest that direct administration of vancomycin to an infected hip joint may not have a detrimental effect on the viability of the MSC population residing in the affected bone tissue and may, therefore, spare the surrounding bone tissue, or at least preserve the capacity for tissue regeneration. The mild anti-osteogenic effects of vancomycin in differentiating pfMSCs may cause local short-term loss of bone mineral density at vancomycin-treated sites. However, we hypothesize that the absence of cytotoxicity of vancomycin on undifferentiated pfMSCs may result in the population of pfMSCs surrounding the affected area remaining intact, which will contribute to repair of the bone tissue and successful osseointegration of the prosthesis upon cessation of vancomycin treatment.

## ACKNOWLEDGMENTS

The research was funded by the National Research Foundation, South Africa. Grants were allocated to H Sadie-Van Gijsen and LMT Dicks. The funders had no role in study design, data collection and interpretation, or the decision to submit the work for publication.

## REFERENCES

1. Kujala UM, Kaprio J, Sarna S. 1994. Osteoarthritis of weight bearing joints of lower limbs in former élite male athletes. Br Med J 308:231–234.

2. Crowninshield RD, Johnston RC, Andrews JG, Brand RA. 1978. A biomechanical investigation of the human hip. J Biomech 11:75–85.

3. Sato T, Sato N. 2015. Clinical relevance of the hip joint: Part I - Review of the anatomy of the hip joint. Int Musculoskelet Med 37:141–145.

4. Zhang Y, Jordan JM. 2010. Epidemiology of Osteoarthritis. Clin Geriatr Med 26:355–369.

5. Song Z, Borgwardt L, Høiby N, Wu H, Sørensen TS, Borgwardt A. 2013. Prosthesis infections after orthopedic joint replacement: the possible role of bacterial biofilms. Orthop Rev (Pavia) 5:65–71.

6. Siopack JS, Jergesen HE. 1995. Total hip arthroplasty. West J Med 162:243–249.

7. Knight SR, Aujla R, Biswas SP. 2011. Total Hip Arthroplasty - over 100 years of operative history. Orthop Rev (Pavia) 3:72–74.

8. Pivec R, Johnson AJ, Mears SC, Mont MA. 2012. Hip arthroplasty. Lancet 380:1768–1777.

9. Lentino JR. 2003. Prosthetic joint infections: bane of orthopedists, challenge for infectious disease specialists. Clin Infect Dis 36:1157–1161.

10. Tande AJ, Patel R. 2014. Prosthetic joint infection. Clin Microbiol Rev 27:302–345.

11. Høiby N, Bjarnsholt T, Givskov M, Molin S, Ciofu O. 2010. Antibiotic resistance of bacterial biofilms. Int J Antimicrob Agents 35:322–332.

12. Pulido L, Ghanem E, Joshi A, Purtill JJ, Parvizi J. 2008. Periprosthetic Joint Infection: the incidence, timing, and predisposing factors. Clin Orthop Relat Res 466:1710–1715.

13. Parvizi J, Aggarwal V, Rasouli M. 2013. Periprosthetic joint infection: Current concept. Indian J Orthop 47:10–17.

14. Wolf M, Clar H, Friesenbichler J, Schwantzer G, Bernhardt G, Gruber G, Glehr M, Leithner A, Sadoghi P. 2014. Prosthetic joint infection following total hip replacement: results of one-stage versus two-stage exchange. Int Orthop 38:1363–1368.

15. Siljander MP, Sobh AH, Baker KC, Baker EA, Kaplan LM. 2018. Multidrug-resistant organisms in the setting of periprosthetic joint infection - diagnosis, prevention, and treatment. J Arthroplasty 33:185–194.

16. Ravi S, Zhu M, Luey C, Young SW. 2016. Antibiotic resistance in early periprosthetic joint infection. Orthop Surg 86:1014–1018.

17. Parvizi J, Alijanipour P, Barberi EF, Hickok NJ, Phillips SK, Shapiro IM, Schwarz EM, Stevens MH, Wang Y, Shirtliff ME. 2015. Novel developments in the prevention, diagnosis, and treatment of periprosthetic joint infections. J Am Acad Orthop Surg 23:32–43.

18. Kirby A, Graham R, Williams NJ, Wootton M, Broughton CM, Alanazi M, Anson J, Neal TJ, Parry CM. 2010. Staphylococcus aureus with reduced glycopeptide susceptibility in Liverpool, UK. J Antimicrob Chemother 65:721–724.

19. Gu H, Ho PL, Tong E, Wang L, Xu B. 2003. Presenting vancomycin on nanoparticles to enhance antimicrobial activities. Nano Lett 3:1261–1263.

20. Zakeri-milani P, Loveymi BD, Jelvehgari M, Valizadeh H. 2013. The characteristics and improved intestinal permeability of vancomycin PLGA-nanoparticles as colloidal drug delivery system. Colloids Surfaces B Biointerfaces 103:174–181.

21. Lotfipour F, Abdollahi S, Jelvehgari M, Valizadeh H, Hassan M, Milani M. 2013. Study of antimicrobial effects of vancomycin loaded PLGA nanoparticles against Enterococcus clinical isolates. Drug Res (Stuttg) 64:348–352.

22. Song C, Guo Z, Ma Q, Chen Z, Liu Z, Jia H, Dang G. 2003. Simvastatin induces osteoblastic differentiation and inhibits adipocytic differentiation in mouse bone marrow stromal cells. Biochem Biophys Res Commun 308:458–462.

23. Heim M, Frank O, Kampmann G, Sochocky N, Pennimpede T, Fuchs P, Hunziker W, Weber P, Martin I, Bendik I. 2004. The Phytoestrogen Genistein Enhances Osteogenesis and Represses Adipogenic Differentiation of Human Primary Bone Marrow Stromal Cells. Endocrinology 145:848–859.

24. Duque G, Rivas D. 2007. Alendronate has an anabolic effect on bone through the differentiation of mesenchymal stem cells. J Bone Miner Res 22:1603–1611.

25. Jacobs FA, Sadie-Van Gijsen H, van de Vyver M, Ferris WF. 2016. Vanadate impedes adipogenesis in mesenchymal stem cells derived from different depots within bone. Front Endocrinol (Lausanne) 7:1–12.

26. Andrews JM. 2001. Determination of minimum inhibitory concentrations. J Antimicrob Chemother 48:5–16.

27. Prakash V, Lewis JS, Jorgensen JH. 2008. Vancomycin MICs for methicillin-resistant Staphylococcus aureus isolates differ based upon the susceptibility test method used. Antimicrob Agents Chemother 52:4528.

28. Wang G, Hindler JF, Ward KW, Bruckner DA. 2006. Increased vancomycin MICs for Staphylococcus aureus clinical isolates from a university hospital during a 5-year period. J Clin Microbiol 44:3883–3886.

29. Fair RJ, Tor Y. 2014. Antibiotics and bacterial resistance in the 21st century. Perspect Medicin Chem 6:25–64.

